# Clearing the FoG: Antifungal tolerance is a subpopulation effect that is distinct from resistance and is associated with persistent candidemia

**DOI:** 10.1101/206359

**Authors:** Alexander Rosenberg, Iuliana V. Ene, Maayan Bibi, Shiri Zakin, Ella Shtifman Segal, Naomi Ziv, Alon M. Dahan, Arnaldo L. Colombo, Richard J. Bennett, Judith Berman

## Abstract

Drug susceptibility, defined by the minimal inhibitory concentration (MIC), often does not predict whether fungal infections will respond to therapy in the clinic. Tolerance at supra-MIC antifungal drug concentrations is rarely quantified and current clinical recommendations suggest it be ignored. Here, we measured and characterized drug-response variables that could influence the outcomes of fungal infections and be generalizable across major clades of *Candida albicans*, one of the most frequently isolated human fungal pathogens. We quantified antifungal tolerance as the fraction of growth (FoG) above the MIC and found that it is clearly distinct from susceptibility/resistance measured as MIC. Instead, tolerance is due to the slow growth of subpopulations of cells that overcome drug stress more efficiently than the rest of the population, and correlates inversely with the accumulation of intracellular drug. Importantly, many adjuvant drugs used together with fluconazole, a fungistatic drug, reduce tolerance without affecting resistance. These include inhibitors of major stress response hubs such as Hsp90, calcineurin, PKC1 and TOR. Accordingly, in an invertebrate infection model, adjuvant combination therapy was significantly more effective than fluconazole alone in treating highly tolerant isolates and did not improve the treatment of isolates with low tolerance levels. Furthermore, isolates recovered from immunocompetent patients with persistent candidemia displayed significantly higher tolerance than isolates that were readily cleared by fluconazole. Thus, tolerance correlates with the response to fluconazole therapy in patients and may help predict whether infections will respond to fluconazole alone. Similarly, measuring tolerance may provide a useful clinical parameter for choosing appropriate therapeutic strategies to overcome persistent clinical candidemia.

## Introduction

A goal of antimicrobial susceptibility testing is to predict the clinical success or failure of antibiotic therapy. Some infections are recalcitrant to drug treatment due to ‘resistance’, which refers to microbial growth in the presence of drug concentrations that inhibit susceptible isolates^1,2^. Susceptibility is commonly measured as the Minimal Inhibitory Concentration (MIC) after 24 h of growth in the presence of drug^3,4^. Fungal infections generally follow the “90/60” rule for predicting therapeutic outcomes based on *in vitro* susceptibility testing: ∼90% of susceptible isolates and 60% of resistant isolates respond to therapy^5-9^. This implies that infection outcomes are influenced by host factors as well as features of the pathogen that are not reflected by the MIC^10,11^. In effect, organisms that cause persistent infections, defined as those that are not cleared by a course of antifungal treatment, have similar susceptibilities to organisms that are readily cleared by a course of antifungal treatment^12^. Accordingly, it is important to identify measurable parameters that can contribute to disease severity.

Only four classes of antifungals are currently in clinical use, and resistance to azoles, including fluconazole (FLC), the most commonly administered antifungal against *Candida* species, is an increasing problem. Altered drug uptake/drug efflux and changes in ergosterol biosynthesis (the target for azole drugs)^13^ are the major known mechanisms of azole resistance. Stress responses have been proposed as a third mechanism of antifungal resistance^14^. A broad range of small molecules enhance antifungal activity *in vitro* and *in vivo*, with inhibitors of Hsp90, calcineurin and TOR the most prominent among them^15-18^. Combination therapy using antifungals together with such inhibitors has been proposed as a promising strategy to extend the efficacy of current drugs^19-24,25,26,27^. In addition, several psychotherapeutic agents such as fluoxetine, fluphenazine or sertraline can enhance FLC activity against fungal species^21,28-30^. Whether such adjuvants affect therapeutic outcomes remains to be addressed.

Persistent candidemia, defined as the failure to clear a bloodstream infection caused by a susceptible organism^12,31,32^, is associated with increased mortality. In one study, the mortality rate was 54% among infections with persistent candidemia and only 3% among those with non-persistent candidemia^33^. Mechanisms underlying persistent candidemia may include variability in the pharmacology of the drug, suboptimal dosing, presence of fungal biofilms on indwelling catheters, and reduced immunity. We posit that some responses to the drug are not captured by measuring the MIC alone and that additional parameters could be used to predict the likelihood that a clinical isolate might respond poorly to antifungal drugs. Furthermore, understanding the mechanisms that underlie these parameters is critical for the development of new therapeutic approaches against persistent *Candida* infections.

MIC measurements have been optimized to minimize or ignore residual fungal growth^9,34,35^, termed ‘tolerance’ or ‘trailing growth’, which has been discussed in the literature for over 20 years^36,37^ and is detected in 25-60% of clinical isolates^4,10,19,38-44^. This recommendation is based upon studies of acute infections using the mouse model of bloodstream candidiasis^45-47^, and the observation that isolates with high trailing growth in mucosal infections respond positively to short term antifungal treatment, despite later recurrence of infection^48^. Trailing growth is sensitive to environmental conditions, including pH, temperature and nutrients^3,49,50^ and is usually detected in liquid cultures. Definitions of ‘tolerance’ vary and generally describe survival or growth above inhibitory concentrations^2,10,28,38,51-56^, detected as slow growth within the zone of inhibition using E-strips or disk diffusion assays ^57,58^, or in broth microdilution assays^38,39^. Tolerance can be affected by several adjuvant drugs^19-24^, iron levels^19,59^, genes involved in vacuolar protein sorting^38,47^, as well as calcium flux^28,55,59-61^. However, the precise relationship between the outcome of fungal infections and tolerance or trailing growth has not been determined. Because tolerant cells continue to divide in the presence of antifungals, we postulated that they contribute to the persistence and/or recurrence of fungal infections.

Here, we measured susceptibility and tolerance in a broad range of clinical isolates spanning the major *C. albicans* clades^62-64^ and asked how these parameters influence fungal infection outcomes. We analyzed disk diffusion assays using *diskImageR*, and quantified the radius (RAD) of the zone of inhibition, a parameter that relates to the MIC^65^, and the fraction of growth (FoG) within the zone of inhibition, a parameter that measures tolerance^19,38,50,66-68^. We found that a range of adjuvant drugs, used in combination with FLC, increased drug cidality by reducing FoG and not MIC and were effective both *in vitro* and *in vivo* at inhibiting strains with high (and not low) FoG levels. Finally, highly tolerant isolates were associated with persistent candidemia, suggesting that knowing the tolerance level of an infecting isolate may have important clinical implications and may inform treatment options.

## Results

### Tolerance measured as FoG is distinct from drug resistance

We quantified drug responses in *C. albicans* isolates from different genetic backgrounds and types of infections using *diskImageR,* a quantitative analysis tool that measures RAD, the radius of the zone of inhibition, an indicator of susceptibility that relates to MIC^69,70^, and FoG, the fraction of growth within the zone of inhibition, relative to the maximum possible growth^65^ (Fig. 1a). A screen of 219 clinical isolates (Supplementary Table 1) revealed that FoG levels ranged widely from 0.10 to 0.85 and did not correlate with RAD levels (Fig. 1b,c), indicating that FoG and RAD measure independent drug responses. FoG was detected in response to other drugs including fungistatic antifungals (azoles, 5-fluorocytosine) and, to a lesser degree, to fungicidal agents (echinocandins, polyenes) (Supplementary Fig. 1a). Antifungal responses varied widely in different *Candida* species as well as in *S. cerevisiae* (Supplementary Fig. 1b). For example, *C. glabrata* exhibited the highest tolerance to azoles, followed by *C. tropicalis* and *C. krusei,* while *S. cerevisiae* had low tolerance to azoles and intermediate tolerance to echinocandins and polyenes.

**Figure 1.**
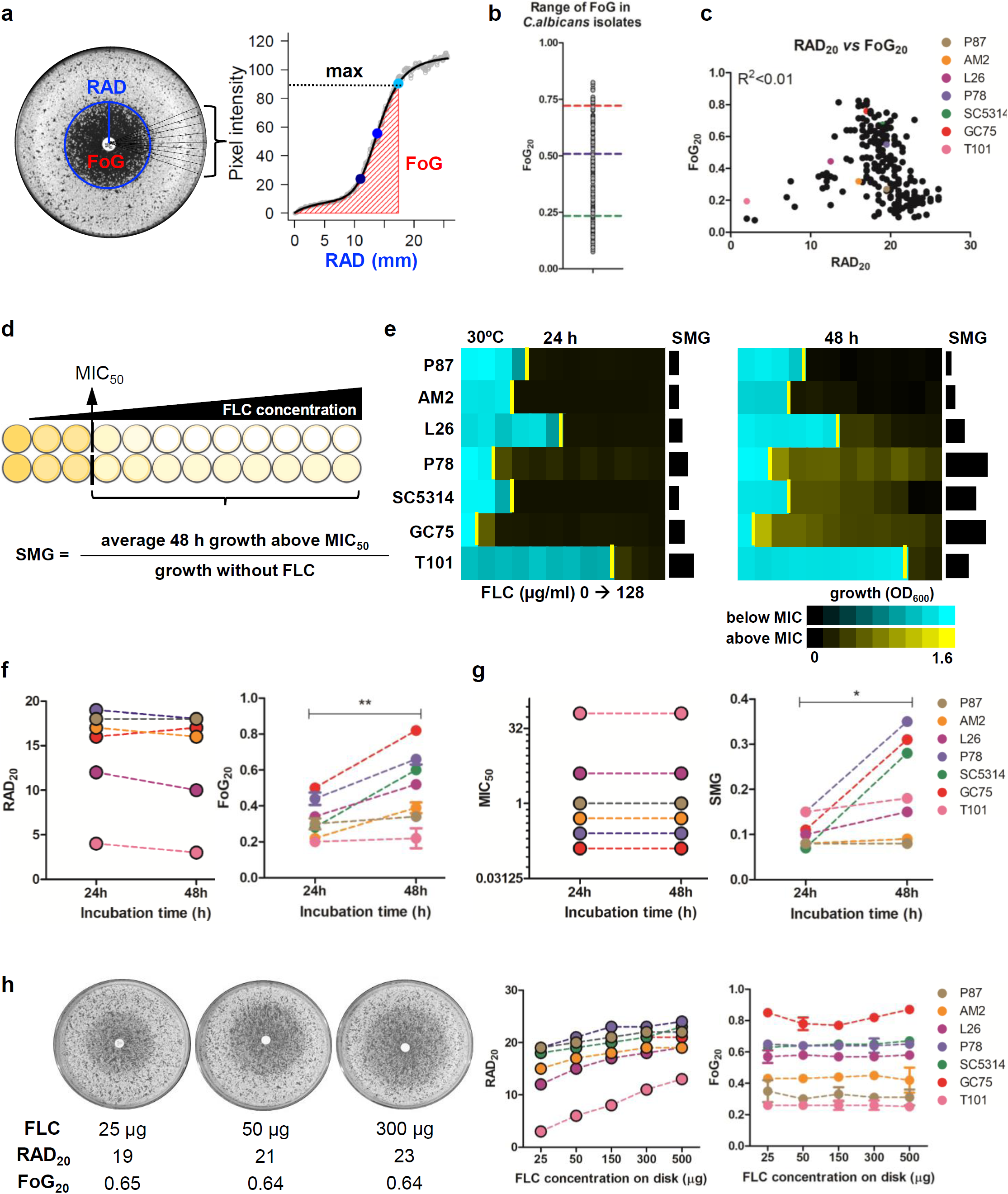
Measuring drug responses of *C. albicans* clinical isolates in disk diffusion assays (DDAs) and in liquid broth microdilution assays (BMDAs). (a) *diskImageR* analysis measures pixel intensity corresponding to cell density, along 72 radii every 5°. The average radius (RAD) represents the distance in mm corresponding to the point where 20%, 50% or 80% growth reduction occurs (light, medium, dark blue dots). The fraction of growth inside the zone of inhibition (FoG) is the area under the curve (red) at the RAD threshold, divided by the maximum area. (b) Range of FoG levels in 219 *C. albicans* clinical isolates. Red, blue and green lines estimate high, medium and low FoG levels, respectively. Unless otherwise specified, disk diffusion assays were performed using a single 25 μg FLC disk and analyzed after 48 h at 30°C. (c) Comparison of FoG_20_ and RAD_20_ for the 219 isolates. (d) Illustration of MIC and supra-MIC growth (SMG) calculations. MIC_50_ was calculated at 24 h as the FLC concentration at which 50% of the growth was inhibited, relative to growth in the absence of drug. SMG was calculated as the average growth per well above the MIC divided by the level of growth without drug. FLC was used in two-fold dilutions (0, 0.125 to 128 µg/ml). (e) Heatmaps illustrating OD_600_ levels for concentrations below the MIC (yellow bar) in cyan and above the MIC in yellow for the seven *C. albicans* isolates from Fig. 1c. Maps show OD_600_ values at 24 and 48 h. (f-g) Effect of incubation time on RAD/MIC and FoG/SMG values. *diskImageR* analysis (f) and corresponding MIC and SMG levels (g) measured at 24 and 48 h for strains as in (d). Of note, truly drug-resistant strains such as T101 (MIC = 64, Fig. 1e), the small zone of inhibition makes FoG measurements less accurate than SMG levels. (h) RAD is concentration-dependent and FoG is concentration-independent as measured with disks containing increasing concentrations of FLC (25, 50 and 300 μg) for strain SC5314 shown for illustration (left); RAD (middle) and FoG (right) levels for strains as listed.

Broth microdilution assays define the MIC_50_ - the lowest drug concentration that inhibits 50% of growth at 24 h (termed MIC throughout the manuscript). Most reports of ‘trailing growth’ or ‘tolerance’ monitor growth at 48 h^48-50,71^. We quantified supra-MIC growth (SMG) at 48 h (average growth per well above the MIC_50_, normalized to growth without FLC; Fig. 1d). Most isolates exhibited some level of SMG at 48 h, while MIC levels remained unchanged between 24 and 48 h (Fig. 1e-g). Thus, SMG provides a parameter that, like FoG, is distinct from MIC.

Importantly, cells exhibiting FoG in disk assays or SMG in broth assays, are not due to the emergence of drug resistance. Thus, for a given isolate, FoG and SMG levels were reproducible for cells taken from inside or outside the zone of inhibition, or for cells taken from wells above or below the MIC (Supplementary Fig. 2a, b). These cells yield progeny indistinguishable from other cells in the population, and thus are the result of phenotypic heterogeneity rather than genetic alteration. Consequently, tolerance is due to growth heterogeneity in the population and is stable for a given isolate.

**Figure 2.**
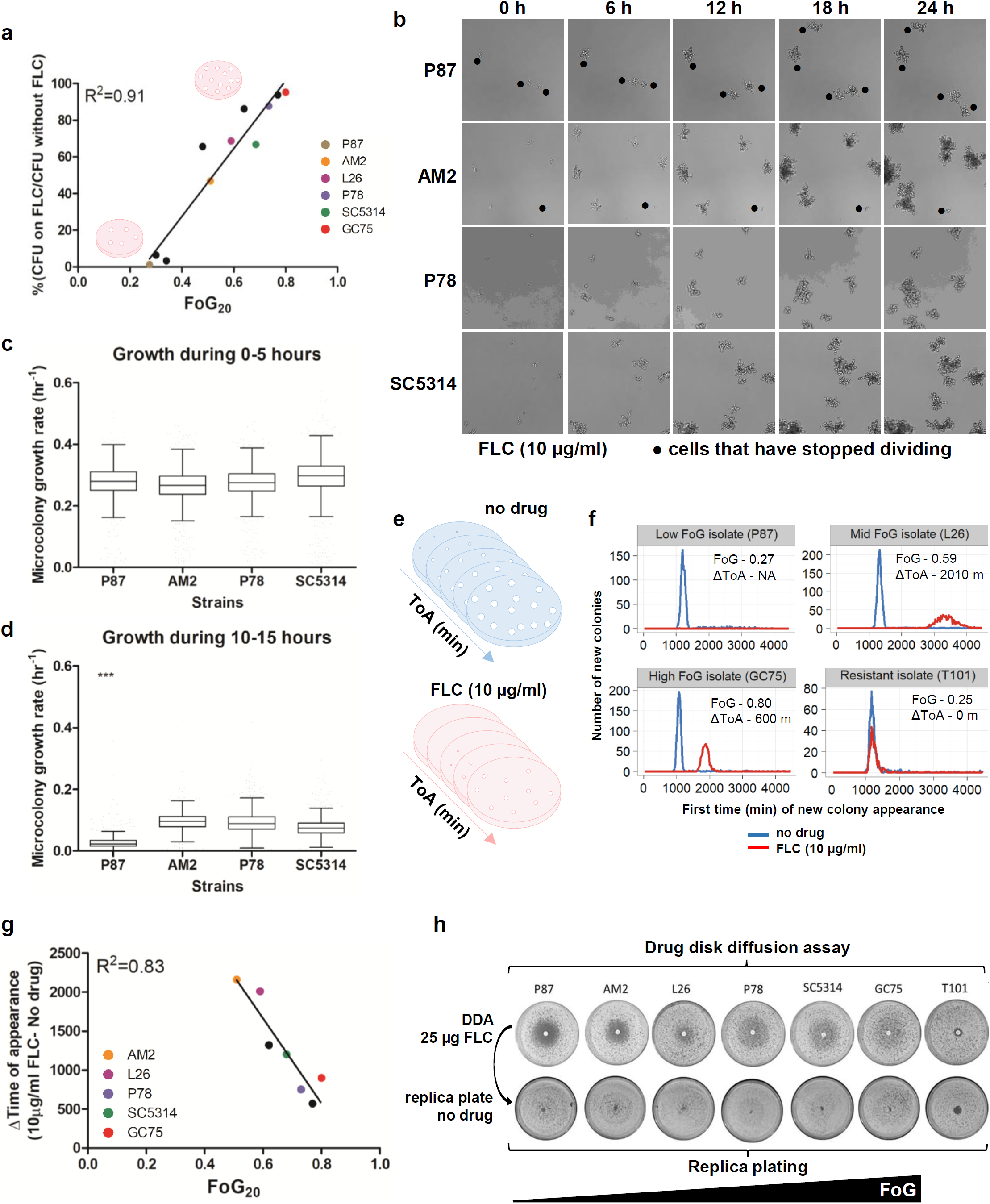
Analysis of cells growing at supra-MIC concentrations. All tested strains, except for T101 (MIC = 64 μg/ml), had FLC MIC values below 10 μg/ml. (a) FoG correlates with the proportion of colonies that grow on 10 μg/ml FLC relative to growth on plates without drug. (b) Microcolony analysis at supra-MIC concentrations of FLC (10 μg/ml). Symbols indicate cells that have stopped dividing in the presence of drug over 24 h. On average, strain AM2 formed fewer microcolonies than strain SC5314, but these were larger than those formed by SC5314. Time lapse videos are available as Supplementary videos 1-4. (c,d) Growth rate analysis of cells growing at supra-MIC FLC (10 μg/ml) during 0-5 h (c) and 10-15 h (d). (e) Schematic of ScanLag analysis^75^ that measures time of colony appearance (ToA), colony growth rate and colony size using desktop scanners. (f) ToA on medium without drug (blue) or with 10 μg/ml FLC (red) for resistant isolate T101, and isolates with different FoG levels. Graphs show the number of colonies (y-axis) at each time point (x-axis). Additional isolates are included in Supplementary Fig. 4. (g) Correlation between FoG_20_ and the difference (Δ) in the ToA of colonies in the presence vs absence of FLC (ΔToA = ToA with FLC – ToA without FLC). (g) Cells that grow within the zone of inhibition are viable, as seen by replica plating of disk diffusion assays grown on casitone without FLC and incubated at 30°C for 48 h.

While SMG and FoG correlated well with each other (R^2^ = 0.82, *P* < 0.01), there was no relationship between FoG and RAD (R^2^ < 0.01), or SMG and MIC (R^2^ = 0.25, *P* < 0.01, Supplementary Fig. 2c). FoG and SMG therefore reflect similar features of growth at supra-MIC concentrations that are distinct from drug susceptibility/resistance, as measured by RAD or MIC, which are concentration-dependent parameters. Accordingly, whereas RAD increased with increasing drug concentrations in the disk (Fig. 1h), FoG remained similar irrespective of drug concentration. Similarly, SMG levels remained constant across a broad range of supra-MIC concentrations (Fig. 1e). Thus, tolerance measured as FoG or SMG is not dependent upon drug concentration, highlighting the distinctive nature of tolerance relative to susceptibility/resistance.

### Environmental modulation of drug responses

FoG and SMG levels were reduced at pH 4.5 relative to pH 7, while RAD/MIC levels remained relatively stable (Supplementary Fig. 3a,b). FoG/SMG values also decreased with higher temperatures (37-41°C) while RAD/MIC levels showed less variation with increasing temperature (Supplementary Fig. 3c,d). For some strains, including SC5314, RAD and/or FoG levels varied considerably depending on the growth media (Supplementary Fig. 3e). A series of strains derived from SC5314 by passaging were particularly sensitive to media differences, with lower RAD and higher FoG levels on rich medium (YPD) than on casitone medium (Supplementary Fig. 3f), emphasizing that assays must be performed under consistent conditions to ensure reproducible results. Importantly, differences in genetic background can have a major effect on how *C. albicans* strains respond to environmental conditions.

**Figure 3.**
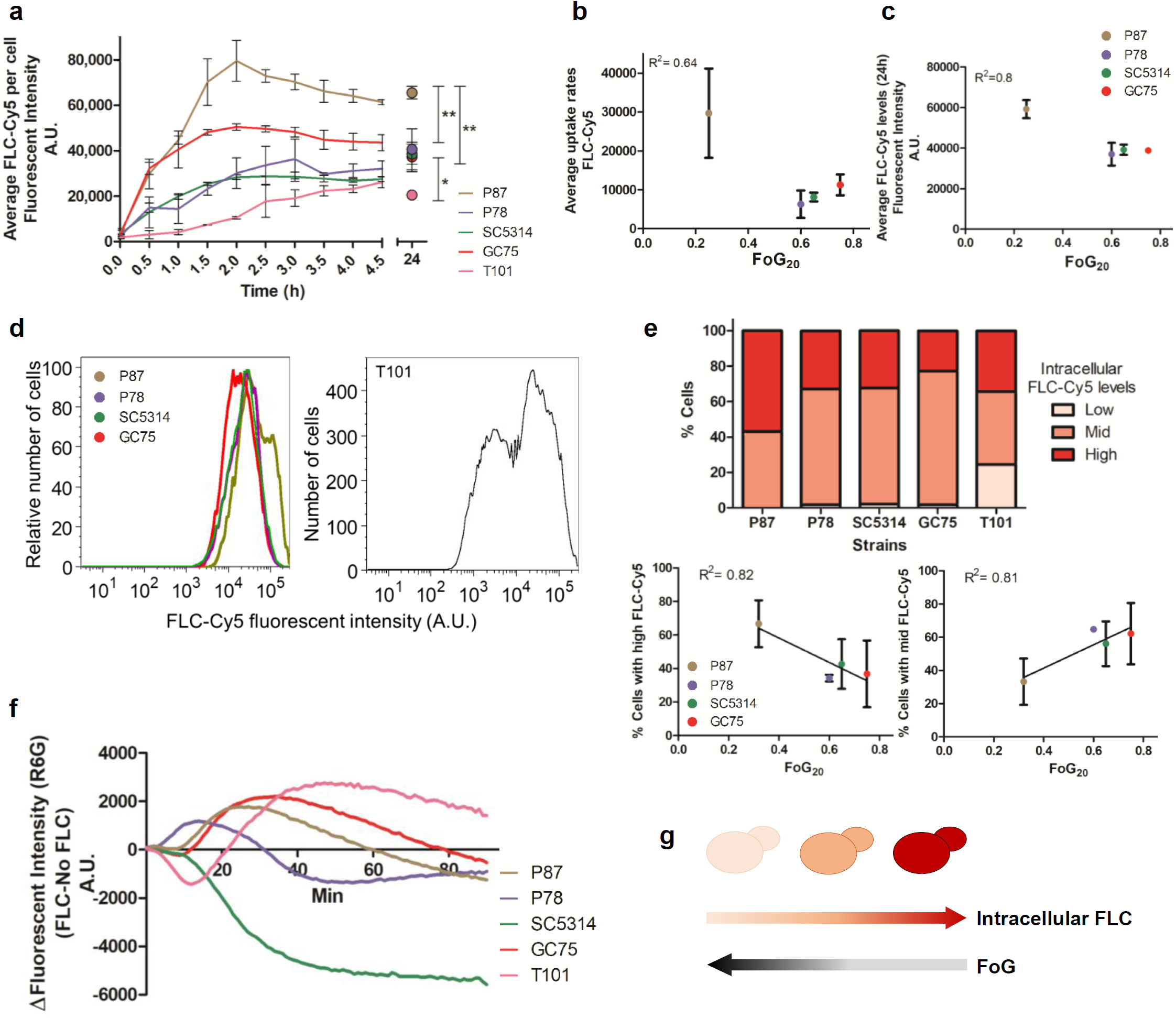
Uptake, efflux and steady state intracellular concentrations of FLC-Cy5. (a) Average intracellular FLC-Cy5 uptake per cell. Uptake curves for 0-5 h and for steady state FLC-Cy5 per cell at 24 h measured by flow cytometry. (b,c) Correlation between FoG_20_ and uptake rate of FLC-Cy5 between 0.5-1.5h (b) and intracellular FLC-Cy5 at 24 h (c). (d) Intracellular FLC-Cy5 fluorescent intensity at 24 h for four strains with similar MIC and diverse FoG levels (left) and for the resistant strain T101 (right) normalized for number of cells (n= 15863 for P87, 15075 of P78, 22406 for SC5314, 18186 of GC75); (e) The proportion of cells (%) with FLC-Cy5 levels at 24 h was divided into thirds: Low (391-422 A.U.), Mid (3912-34643 A.U.) and High (3.5×10^4^-3.14×10^5^ A.U.) (upper panel). Proportion of cells (%) with high (lower left panel) or mid (lower right panel) intracellular FLC-Cy5 concentration at 24 h vs FoG_20_ levels of the strains. (f) Efflux of Rhodamine 6G (R6G) normalized by culture density (OD_600_). Efflux curves represent data from one (of two) experiments and the curves show fluorescence intensity recorded over 90 min and calculated as follows: Δ[FLC(with Glucose - without Glucose) – No drug(with Glucose - without Glucose)]. (g) Cartoon illustrates the steady state drug levels in cells with high to low FoG levels.

### Subpopulation growth dynamics

To ask if tolerance reflects the size of a dividing subpopulation at supra-MIC concentrations, we compared the number of cells that form colonies (CFUs) in the presence of supra-MIC FLC concentrations to total CFUs without drug. CFUs on drug ranged between 2% and 98% of the total population and correlated with FoG levels (Fig. 2a, R^2^ = 0.91, *P* < 0.01). The proportion of the subpopulation that grew at supra-MIC FLC in liquid cultures did not change with inoculum size (Supplementary Fig. 3g), indicating that FoG/SMG is independent of cell density. Importantly, these results establish that FoG levels represent the proportion of the population that can form colonies or grow above the MIC.

Growth dynamics were also monitored using microcolony assays, in which time-lapse microscopy follows colony area/size (Fig. 2b)^72^. As expected^73^, exponentially growing cells plated on 10 μg/ml FLC did not stop growing immediately. Rather, most cells continued dividing for ∼5 h, and then slowed or stopped dividing to different degrees (Fig. 2c,d and Supplementary videos 1-4). In high FoG isolates (e.g., SC5314), a large proportion of the cells yielded microcolonies, while in low FoG isolates (e.g., P87), more cells stopped dividing (black dots in Fig. 2b). Differences in growth rates became apparent after 10-15 h in FLC (Fig. 2d), supporting the idea that different subpopulations have different growth dynamics, and that high FoG strains produce a larger subpopulation of growing colonies than low FoG isolates.

The dynamics of colony growth on agar plates was also examined using *ScanLag,* a flatbed scanner/image analysis pipeline^74^ that we adapted for use with *C. albicans*. Here, the area occupied by light pixels and the change in this area over time are proxies for colony size and growth rate, respectively^74^. Different *C. albicans* isolates exhibited different initial time of colony appearance (ToA) on medium without drug (Supplementary Fig. 4). At 10 μg/ml FLC, the ToA was delayed relative to the drug-free condition (except for resistant strains such as T101, Fig. 2e,f). Notably, the ΔToA (ToA on FLC – ToA without FLC) of high FoG isolates was shorter than that of low FoG isolates (Fig. 2f), and correlated with overall FoG levels (Fig. 2g). This suggests that high FoG strains overcome the inhibitory pressures of the antifungal more efficiently than low FoG strains. Consistent with this, in liquid assays with drug, highly tolerant isolates also displayed shorter lag times than related isolates with low tolerance levels (Supplementary Fig. 5a).

**Figure 4.**
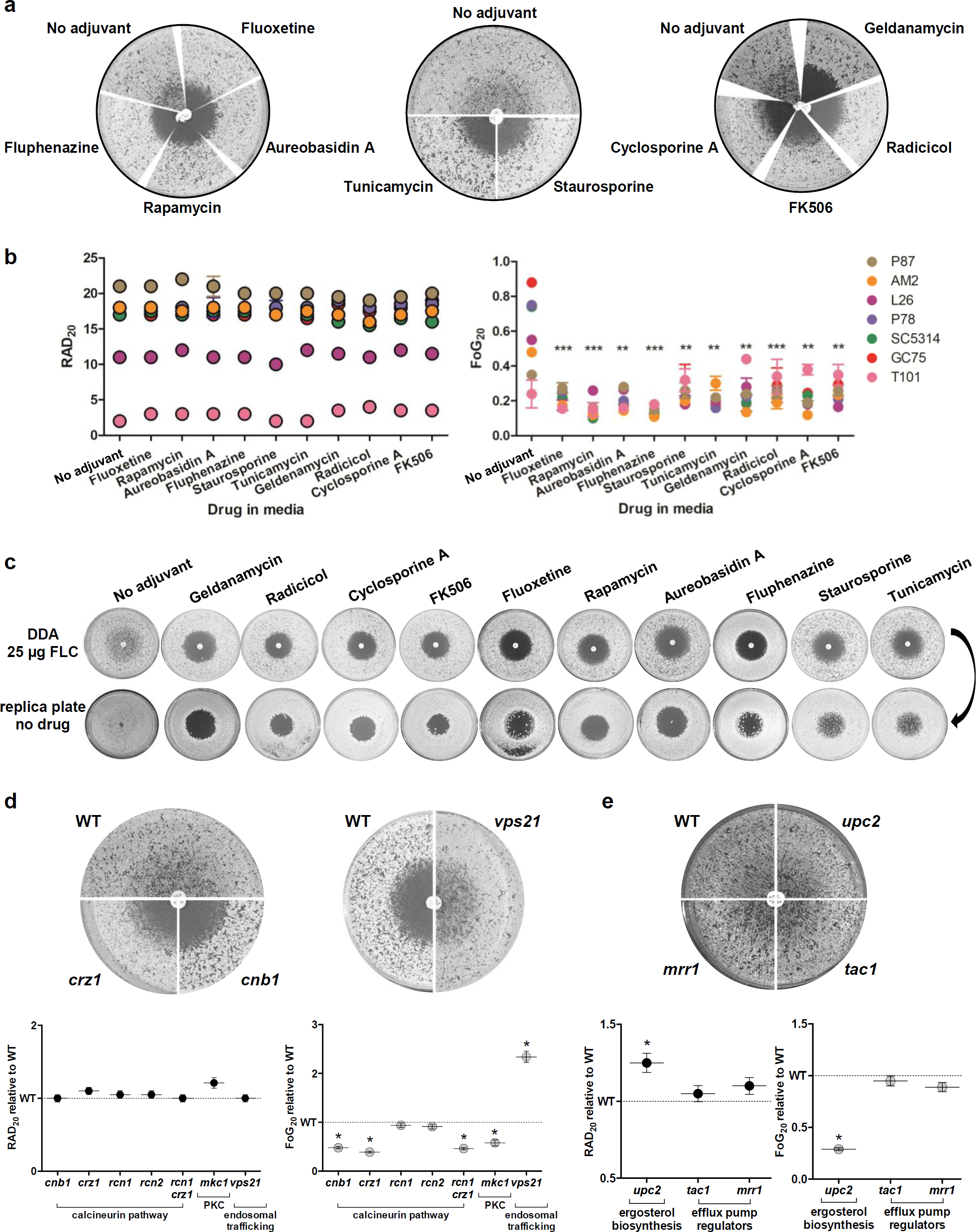
Adjuvant drugs significantly reduce FLC tolerance but not resistance and render FLC cidal. (a) Disk diffusion assays performed with 25 μg FLC on casitone plates supplemented with adjuvant drugs 20 μg/ml fluoxetine, 5 ng/ml aureobasidin A, 0.5 ng/ml rapamycin, 10 μg/ml fluphenazine, 12.5 ng/ml staurosporine, 0.25 μg/ml Tunicamycin, Hsp90 inhibitors (0.5 μg/ml geldanamycin and 0.5 μg/ml radicicol) and calcineurin inhibitors (0.5 μg/ml FK506 and 0.4 μg/ml cyclosporine A) shown for strain SC5314. (b) RAD and FoG levels performed on disk diffusion assays with FLC and adjuvants. (c) Effect of drug adjuvants and pathways inhibitors on the viability of cells growing inside the zone of inhibition. FLC disk diffusion assays of SC5314 without or with adjuvant were replica plated (after removal of the drug disk) onto casitone plates (without FLC or adjuvants) and incubated at 30°C for 48 h. (d,e) FLC disk diffusion assays performed using a series of mutants carrying deletions in gene encoding the calcineurin subunit Cnb1, the calcineurin responsive transcription factor Crz1, calcineurin regulators Rcn1 and Rcn2, MAP kinase Mkc1, vacuolar trafficking protein Vps21 (d), ergosterol biosynthesis regulator Upc2, and efflux pump regulators Tac1 and Mrr1 (e). These mutants as well as the *rcn1 crz1* double mutant were analyzed by *diskImageR,* and RAD and FoG levels are shown relative to the isogenic parental strains. All pictures in this figure are representative of two biological replicates, asterisks denote significant differences relative to corresponding parental strains, *P* < 0.05.

**Figure 5.**
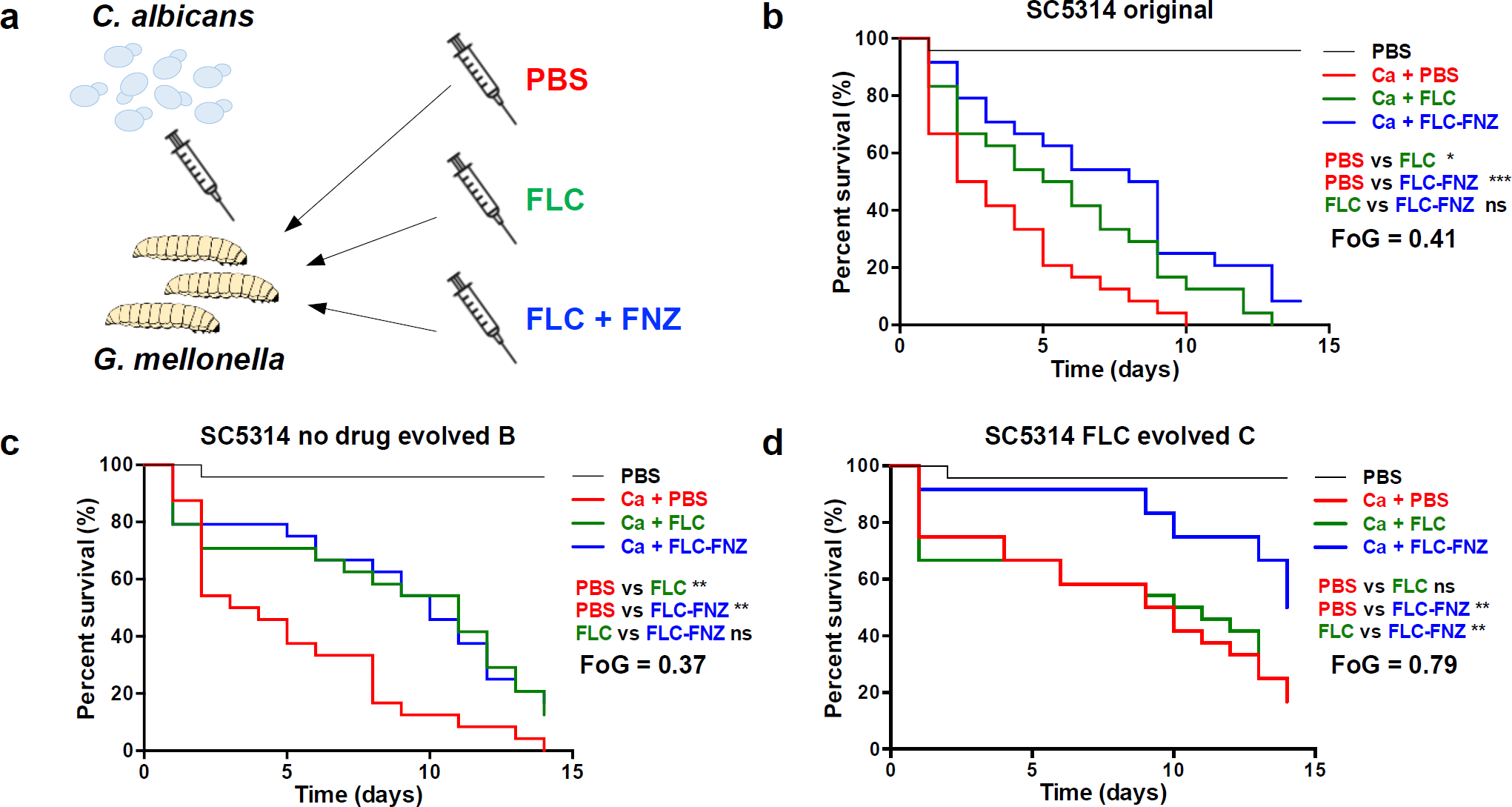
Combination therapy partially rescues systemic infection of *G. mellonella* by high FoG *C. albicans* isolates. (a) *G. mellonella* larvae were injected with 3 × 10^5^ yeast cells/larvae followed by a second injection with either PBS, FLC alone or FLC and FNZ within 90 min after the first injection. (b-d) Survival curves of larvae infected with SC5314, low FoG (c) and high FoG (d) isogenic derivatives. Each curve represents a group of 24 larvae which were monitored daily for survival for up to 14 days after infection. *P* values represent results of log-rank test comparing different treatment conditions with significance values as follows: *, *P* < 0.05; **, *P* < 0.01; ***, *P* < 0.001.

A number of *C. albicans* growth parameters did not correlate with FoG levels. These included cell viability in the presence of drug (Fig. 2h), consistent with FLC being fungistatic rather than fungicidal. Unlike tolerant bacteria, which display reduced growth rates and longer colony appearance times^2,75-77^, antifungal tolerance did not correlate with slower growth in three sets of related *C. albicans* strains grown in liquid without drug, including: (1) passaged derivatives of SC5314 with altered FoG levels, (2) a slow-growing clinical isolate (P37005) and faster growing derivatives^78^ and (3) a *clb4* mutant^79^ with delayed cell cycle progression relative to control strains (Supplementary Fig. 5b). Furthermore, exponential and stationary phases yielded similar FoG levels (Supplementary Fig. 5c), unlike phenotypic drug resistance in bacteria^80^. Growth parameters (colony size and growth rate, liquid growth rate and lag phase duration) also failed to show any correlation with FoG levels (Supplementary Fig 5d,e). This implies that tolerance is not simply a reflection of faster growth in the presence or absence of drug. Rather, it reflects two aspects of *C. albicans* growth dynamics: a larger subpopulation able to form colonies at supra-MIC drug concentrations (Fig. 2a), and a faster relative recovery in response to fungistatic pressures.

### Decreased intracellular drug levels underlie increased tolerance

A recently developed fluorescent azole probe, FLC-Cy5^81^, provides a powerful tool to monitor intracellular drug levels and uptake rates using flow cytometry (Fig. 3a). The initial rate of FLC-Cy5 uptake per cell varied between strains and showed a weak inverse correlation with FoG levels (R^2^ = 0.64, *P* = 0.12, Fig. 3b). By contrast, steady state FLC-Cy5 levels at 24 h correlated inversely with FoG levels (R^2^ = 0.8, *P* < 0.01, Fig. 3c). Cells were categorized as having low, mid, and high levels of FLC-Cy5 (at 24 h). FoG levels inversely correlated with the proportion of cells that contained high FLC-Cy5 levels (R^2^ = 0.82, *P* = 0.04) and positively correlated with the proportion of cells containing mid FLC-Cy5 levels (R^2^=0.81, *P* = 0.05, Fig. 3d,e).

Based on the assumption that the steady state drug levels are influenced by both drug efflux and uptake, we measured efflux of rhodamine-6-G (R6G), a dye removed primarily via the ABC class of transporters^82^. In general, the rate of R6G efflux was higher in the presence, than in the absence, of FLC, with SC5314 as the notable exception (Fig. 3f and Supplementary Fig. 6a). Yet, efflux rates did not correlate with FoG under any of the conditions tested (Supplementary Fig. 6b, R^2^ < 0.2). Thus, the determinants of steady state intracellular FLC levels are complex, and include uptake and efflux rates as well as other factors (Fig. 3g).

**Figure 6.**
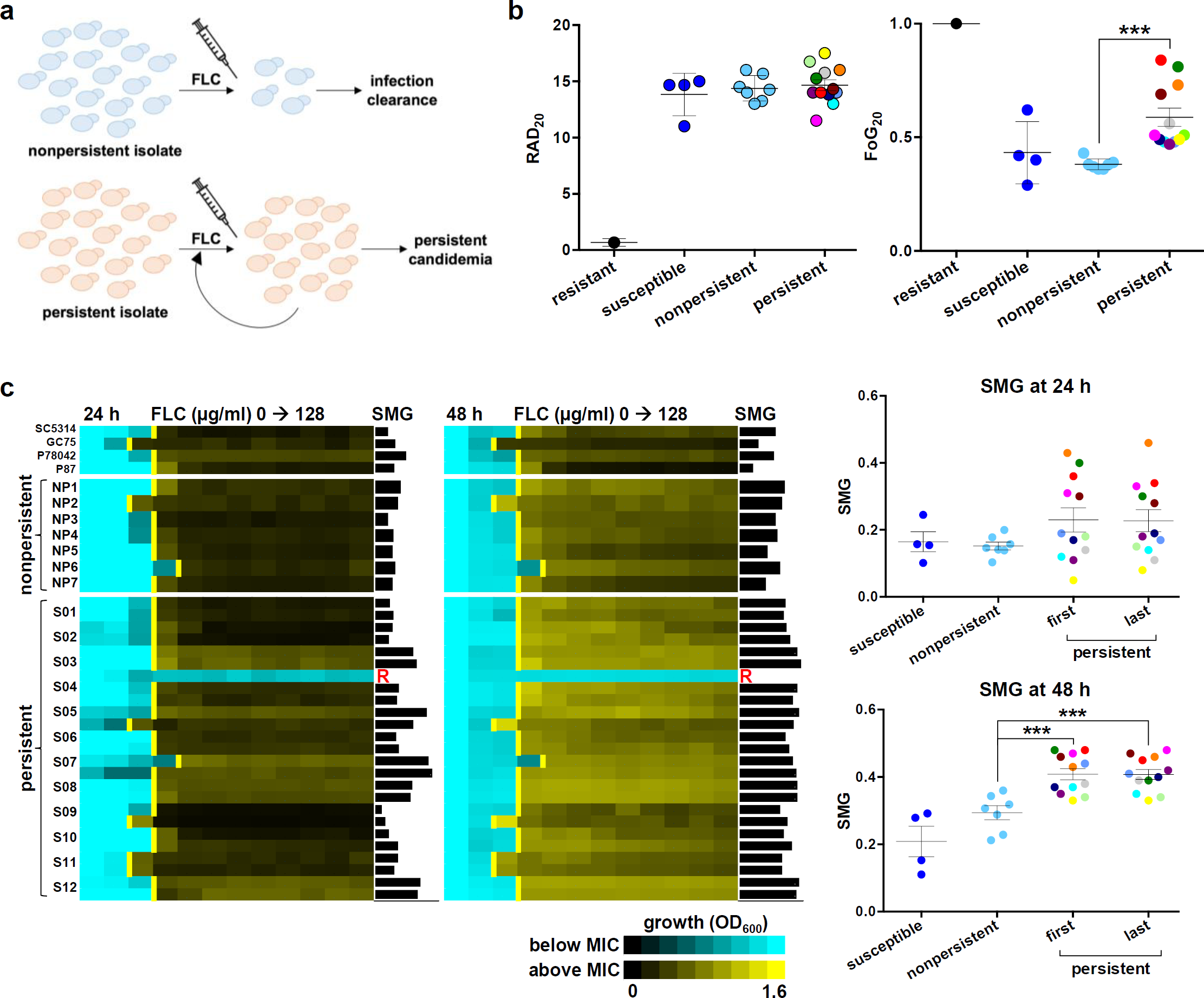
FoG and SMG levels differ between persistent and non-persistent isolates. (a) *C. albicans* isolates from bloodstream infections were either efficiently cleared by a single course of FLC treatment (non-persistent) or persisted in the host despite multiple rounds of FLC therapy. (b) FoG and RAD levels for drug-susceptible (S, n = 4) isolates SC5314, GC75, P78042 and P87, resistant (R, n =1) isolate P60002^31^, non-persistent isolates (NP, n = 7) and the first patient isolate from each of the series of clinically persistent strains (P, n = 12). The final isolate of the S03 series of persistent strains had become FLC-resistant (MIC > 128 μg/ml, Supplementary Fig. 10a), therefore the penultimate isolate, which remained susceptible, was used across analyses. Asterisks indicate significant differences between persistent and non-persistent isolates (t-test, *P* < 0.001). (c) Broth microdilution assays showing MIC and SMG levels at 24 and 48 h for the susceptible control strains, the non-persistent isolates as in (b) and for both the initial and final isolates for each of the 12 clinically persistent series (S01-12). The final isolate in S03 became FLC resistant (R), therefore the penultimate isolate in this series was included as well. Asterisks indicate significant differences between persistent and non-persistent isolates (t-test, *P* < 0.001).

### Adjuvant drug combinations clear tolerance and do not alter resistance

Combination therapies has the potential to extend the lifespan of the few available antifungal drug classes^24,83-85^. We tested a series of adjuvant drugs to determine their quantitative effect on fungal responses when used together with FLC. Most adjuvants significantly reduced FoG, including geldanamycin and radicicol, which inhibit Hsp90 activity^15,27,61,86^, cyclosporine A and FK506, which inhibit calcineurin^87-89^ and staurosporine, which inhibits PKC1 activity^90^. In addition, other adjuvants not directly connected to Hsp90 function also reduced FoG, including aureobasidin A, which inhibits sphingolipid biosynthesis^20^; rapamycin, an inhibitor of the mTOR signaling pathway^22,91^; tunicamycin, an inducer of the unfolded protein response pathway^92-94^; fluoxetine, a serotonin inhibitoR^21^; and fluphenazine, an antipsychotic drug that stimulates ABC transporter expression and indirectly inhibits calcineurin via calmodulin^23,28,95^. FLC + adjuvant cleared tolerance for all 7 clinical isolates tested (Fig. 4a,b), as well as for a longitudinal series from a single HIV patient^96,97^ that had acquired resistance over time, and for a set of persistent and non-persistent clinical isolates (Supplementary Fig. 7a-d). Importantly, most adjuvants had little or no effect on RAD/MIC levels (Fig. 4a,b and Supplementary Fig. 7a-d), and thus are not expected to affect resistance in standard susceptibility assays^34,98^.

Hsp90-dependent responses affected tolerance and not susceptibility/resistance measured as MIC (at 24 h). Consistent with this, the effects of temperature and Hsp90 inhibitors correlated well with FoG (R^2^ = 0.93, Supplementary Fig. 7e) and only partially with RAD (R^2^ = - 0.61, Supplementary Fig. 7f). Thus, temperature inhibition of Hsp90 affects tolerance and not resistance *per se*. Furthermore, FoG levels were not affected by Hsp90 steady state protein levels, as the amount of Hsp90 did not correlate with FoG (R^2^ = 0.07, Supplementary Fig. 7g). Because inhibitors of calcineurin and Hsp90 are known to affect FLC cidality^15,24,27,61,67,88,99,100^, we asked whether other adjuvant drugs have a similar effect. All tested combinations were cidal (Fig. 4c and Supplementary Fig. 7h), indicating they have a profound effect on cell viability at supra-MIC FLC concentrations and implying that stress pathways make essential contributions to tolerance.

Consistent with the adjuvant drug responses, genes involved in several pathways, including calcineurin-^101^ and PKC signaling^90^, primarily affect FoG and not RAD (Fig. 4d, Supplementary Fig. 8, Supplementary Tables 3, 4, and Supplementary Text). In addition, *IRO1*, which is involved in iron homeostasis^19^ and *VPS21,* which is involved in vacuolar trafficking^38^, affected FoG and not RAD. Thus, many genetic pathways affect the ability of subpopulations of cells to grow in the presence of supraMIC FLC concentrations, including Hsp90-dependent and -independent responses that are due to tolerance rather than resistance.

### Adjuvant drugs improve FLC efficacy against high FoG strains *in vivo*

Given that the beneficial effects of adjuvant drugs are more evident on high FoG strains, we asked if adjuvant therapy would distinguish between levels of tolerance in an infection model. *Galleria mellonella* larvae were used to test potential *in vivo* differences between isolates with different tolerance levels, both with respect to their ability to cause disease, and with their ability to respond to therapy with FLC alone or FLC plus the adjuvant fluphenazine^29,102^ (FNZ, Fig. 4a). A series of isolates was derived by passaging the SC5314 strain in the presence or absence of FLC. The resulting isolates had indistinguishable RAD but distinct FoG levels relative to the parental strain (Supplementary Fig. 9a and Supplementary Table 1).

*G. mellonella* larvae infected with fungal cells were treated with either PBS (control), FLC, or combination therapy (FLC and FNZ) (Fig. 5a). SC5314 and derived strains with similar FoG levels killed all larvae within 9-14 days. Importantly, these lower FoG isolates had similar responses to FLC and to the FLC+FNZ combination: they rescued up to 17% of the infected larvae (Fig. 5b,c and Supplementary Fig. 9b). By contrast, high FoG strains displayed delayed killing and decreased virulence in the larvae, and did not respond to FLC alone (Fig. 5d and Supplementary Fig. 9b). Notably, the FLC+FNZ combination significantly improved the outcomes for high FoG strains relative to FLC alone and PBS-control groups: larval death was delayed and up to 50% of larvae infected with high FoG strains survived the experiment (Fig. 5d and Supplementary Fig. 9b). Similar results were obtained with a low FoG clinical isolate (readily cleared by FLC therapy) when compared with a high FoG isolate (that persisted in the bloodstream during multiple rounds of FLC therapy) (Supplementary Fig. 9c). Presumably, FLC+FNZ significantly decreased larval killing by rendering FLC cidal for those cells growing at supra-MIC concentrations. Thus, *in vivo* responses to azoles and to combination therapies differed for different isolates according to their tolerance levels.

### Persistent candidemia is associated with highly tolerant isolates

Echinocandins are considered the optimal line of therapy for patients with candidemia, yet FLC remains a frontline drug for systemic *C. albicans* infections^103^, especially in low income countries^104^. Treatment with FLC often fails despite isolates being drug-susceptible when tested *in vitro,* and rates of persistent candidemia are often higher in patients treated with FLC rather than with echinocandins^105,106^. To examine possible connections between susceptibility and tolerance levels, we collected sets of clinical isolates from patients with candidemia that were either efficiently cleared by a single course of FLC treatment (non-persistent) or that persisted in the host despite extended FLC therapy (Fig. 6a, Supplementary Fig. 10a, see detailed isolate description in Methods).

The drug responses of persistent and non-persistent isolates and several control strains displayed similar RAD and MIC levels (0.25-1 μg/ml), well below those of resistant strains (clinical MIC breakpoint = 4 μg/ml), when assayed at both 24 and 48 h (Fig. 6b and Supplementary Fig. 10b,c). Strikingly, FoG/SMG levels differed significantly between the two groups of clinical isolates, especially when compared at 48 h (Fig. 6b,c) and, as expected, correlated well with one another (Supplementary Fig. 10d, R^2^ = 0.62). Importantly, persistent candidemia isolates displayed higher FoG/SMG levels than those readily cleared upon FLC treatment. In fact, many persistent isolates had FoG levels higher than 0.5, indicating that over half of the cells in the population could grow, albeit slowly, in the presence of FLC (Fig. 6b). Taken together, these experiments demonstrate that tolerance is an intrinsic property distinct from drug susceptibility, and that tolerance levels correlate with infections that resist azole treatment in the clinic.

## Discussion

Currently, decisions about therapeutic strategies to treat *Candida* infections are based upon patient status, infecting species and antifungal susceptibility, with clinical MIC assays designed to avoid the detection of tolerance^3,34^. Here, quantification and characterization of tolerance across *C. albicans* isolates found that many of them exhibit tolerance, defined as the ability of a subpopulation of cells to grow slowly at supra-MIC concentrations. Tolerance is related to phenomena previously described as ‘trailing growth’ ^3,38,39,49,50,67,71,107^, ‘tolerance’^10,19,40-42^ or Hsp90-dependent ‘resistance’^15,51,61,90,108^. Studying growth dynamics and quantifying tolerance revealed that it is: 1) a subpopulation effect and the size of the subpopulation is stable; 2) due to slow growth in drug stress; 3) correlates with intracellular drug levels; 4) dependent upon stress response pathways, and 5) mechanistically distinct from resistance. Several adjuvants completely cleared FoG without altering susceptibility (MIC) and, when combined with the fungistatic drug FLC, yielded a fungicidal cocktail. Consistent with this, combination therapy was most effective *in vivo* at treating infections by high tolerance strains, whereas the adjuvant did not improve FLC efficacy against low FoG isolates. Furthermore, *C. albicans* isolates that cause persistent candidemia exhibited significantly higher tolerance than isolates readily cleared by FLC. Thus, quantitatively measuring tolerance levels of infecting isolates may provide important prognostic insights concerning both the success of FLC therapy as well as the potential efficacy of combination therapies.

The subpopulation effect of tolerance is readily detected on agar plate assays, where individual cells and their clonal progeny are evident; in liquid assays, cells become mixed and subpopulation effects are more difficult to distinguish. Growth dynamics in colonies^74^, microcolony formation^73^ and liquid growth assays all suggest tolerance correlates with the degree of growth after drug exposure. This implies that a subpopulation of cells is inherently more able to adapt to drug than the rest of the population. Since the size of such subpopulations appears stable, we suggest that tolerance is a function of cell physiology and environmental responsiveness that differs between genetic backgrounds and growth conditions.

We discovered that isolates with higher tolerance levels have more cells with lower intracellular drug levels, but tolerance is not a function of either uptake or efflux rates, despite the well-known role of efflux in drug resistance^109-118^. The factors other than uptake and efflux that impact intracellular FLC levels remain to be determined.

### Adjuvant drugs increase FLC cidality and efficacy via tolerance, not resistance

A novel insight from this work is that inhibitors of cellular stress improve fluconazole efficacy, primarily via their effect on tolerance and not resistance. This has two important implications. First, the subpopulation nature of tolerance suggests that these stress pathways must exhibit cell-to-cell variation. Second, the contributions of stress response pathways differ considerably between isolates, albeit in a manner that maintains population heterogeneity (and the size of the tolerant subpopulation) as a heritable feature of the given strain.

Tolerance is clearly related to the Hsp90-dependent response and distinct from *bona fide* resistance. Hsp90 and calcineurin are well known to affect fungal drug responses, including the cidality of azoles^19,67,87,89,119-122^. How the TOR pathway, the unfolded protein response, PKC signaling, sphingolipid synthesis, iron homeostasis, as well as fluoxetine and fluphenazine affect drug responses and increase FLC cidality remains to be determined.

The distinction between resistance and tolerance is consistent with the mechanisms that impact them: resistance mechanisms directly affect the drug target or its concentration in the cell, thereby enabling efficient growth in the presence of the drug^14,111^. By contrast, tolerance reflects stress response strategies that are indirect and may enable survival despite the continued ability of the drug to interact with its target, to remain in the cell and to affect cell growth. Stress responses can modulate physiology, for example membrane^59,123,123^ or cell wall integrity^124^ and the aggregation of proteins into stress granules or other physiological switches^126^, thereby minimizing deleterious cellular responses to the drug.

### Differences between antifungal and antibacterial tolerance

The growth properties of cells at supra-MIC concentrations measured here with FLC, a fungistatic drug, contrast with those measured for antibacterial tolerance in bactericidal drugs. This likely reflects distinct modes of action between static and cidal drugs as well as different molecular mechanisms of stress responses in eukaryotes and bacteria. Antibacterial tolerance generally involves a longer lag phase for the majority of the population^75-77^. By contrast, antifungal tolerance involves the earlier appearance of a subpopulation of cells that continue to grow in the presence of drug and, unlike ‘phenotypic resistance’ described for bacteria^127^, is not dependent upon growth phase or cell density.

### Clinical implications

This study revisited the question of whether supra-MIC growth is clinically relevant and, by inference, whether the stress pathways that mediate it represent important drug targets. Previous work suggested that trailing growth was not important for virulence in either mouse models^45,46^ or infection outcomes in the human host^48^. However, these studies analyzed short-term responses, rather than persistence or recurrence of *Candida* infection over longer time frames. Here, both *in vivo* and clinical studies provide a proof of principle suggesting that retrospective and prospective clinical studies that measure tolerance are warranted. We posit that monitoring FoG/SMG in standard clinical assays may have prognostic value for the likelihood of a persistent infection, for the responsiveness of a given isolate to FLC, as well as for the synergism of an adjuvant drug with FLC. Furthermore, in developing countries where azoles remain the major antifungals in use, measuring tolerance may be especially relevant and the addition of low cost adjuvant drugs could significantly impact treatment outcomes (e.g.,^128^). Understanding the contribution of tolerance to the progression of fungal infections not only can provide fundamental insights into the biology of fungal subpopulation behaviors, but also has the potential to inform clinical practices.

## Methods

### *C. albicans* isolates

All strains (Supplementary Table 1) were streaked onto rich media (YPD) plates and grown for 24 h at 30°C. A single colony from each strain was arbitrarily selected and frozen in 15% glycerol and stored at −80°C for all assays. For each mutant tested, we used genetically matched parental control strains and, where appropriate, nutrient supplements were used to compensate for auxotrophies (Supplementary Table 4).

<u>Isogenic SC5314 low and high FoG isolates</u> were obtained by sequentially passaging SC5314 every 24 h in YPD (1/100 dilutions, ∼84 generations) either with or without 1 μg/ml FLC. <u>Persistent and non-persistent clinical isolates</u> were obtained from patients with candidemia that were investigated by Dr Colombo during several candidemia surveys^129^. Fungal isolates were obtained from two groups of patients: patients with a single episode of candidemia, for which infection was resolved after the first course of antifungal therapy, and patients with persistent candidemia. Persistent candidemia was defined here as two or more blood cultures positive for *C. albicans*, on one or more days apart, despite at least 3 days of antifungal therapy with FLC. Non-persistent isolates (NP1-7) were cleared from the bloodstream soon after FLC treatment was initiated and treatment was continued for an average of 16 days. In contrast, persistent infections in 12 patients yielded serial isolates both prior to and throughout the course of treatment, with 3 to 9 isolates per patient treated for an average of 20 days (P, S01-S12, Supplementary Fig. 10a). Persistent candidemia in these patients occurred despite clinical MIC assays having established that all isolates were FLC susceptible, with MIC levels < 1 μg/ml. While each patient had distinct clinical backgrounds and trajectories, the two groups were matched in terms of age, underlying conditions, time of central venous catheter removal, and first line of antifungal therapy. Patients received intravenous FLC as first line of therapy and the initial time of therapy did not differ between persistent and non-persistent infections. In some cases of persistent candidemia, antifungal therapy was continued with either caspofungin or amphotericin B (Supplementary Fig. 10a). Of the 19 patients, 7 patients died during the 30 day follow up, 6 of which were unable to clear the fungal infection, although causality between infection and death could not be determined. Isolates from patients with cancer, neutropenia, endocarditis, deep-seated *Candida* infections, corticosteroid or other immunosuppressive drug exposure were excluded from the study. All ethical regulations were observed and the study was approved by the Ethical Committee of the Federal University of São Paulo (January, 2016, NO 9348151215).

### Strain construction

To disrupt the two alleles of the *RCN2* gene, we amplified (using primers BP1440 and BP1441, Supplementary Table 5) the flanking sequence of ORF C7_01700W using BJB-T 2 (*HIS1*) and BJB-T 61 (*ARG4*) plasmids as a template (Supplementary Table 5). Strain YJB-T65 was transformed with this PCR product to generate heterozygote mutant YJB-T 2214 (*rcn2*::*HIS1*) and subsequently homozygote mutant YJB-T2227 (*rcn2::HIS1/rcn2*::*ARG4*). The disruptions were verified using primers BP1444 and BP1445.

### Broth micro dilution assays

Minimal inhibitory concentration (MIC) for each strain was measured using CLSI M27-A2 guidelines^98^ with minor modifications as follows. Strains were streaked from glycerol stock onto YPD agar and grown for 24 h at 30°C. Colonies were suspended in 1 ml PBS and diluted to 10^3^ cells/ml in a 96-well plate with casitone containing a gradient of two-fold dilutions per step of FLC, with the first well contain no drug. For persistent and non-persistent clinical isolates, cells were grown overnight in YPD at 30°C and diluted to 10^4^ cells/ml in YPD containing a gradient of two-fold dilutions per step of FLC. MIC_50_ levels were determined after 24 h or 48 h by taking an optical density reading (OD_600_) by a Tecan plate reader (Infinite F200 PRO, Tecan, Switzerland). MIC_50_ levels (shown as yellow lines on broth microdilution assays heatmaps) were determined as the point at which the OD_600_ had been reduced by ≥50% compared to the no-drug wells.

### Disk diffusion assays

The CLSI M44-A2 guidelines for antifungal disk diffusion susceptibility testing^130^ were followed with slight modifications. In brief, strains were streaked from glycerol stocks onto YPD agar and incubated for 24 h at 30°C. Colonies were suspended in 1 ml PBS and diluted to 1 × 10^6^ cells/ml. 2 × 10^5^ cells were spread onto 15 ml casitone plates (9 g/l Bacto casitone, 5 g/l yeast extract, 15 g/l Bacto agar, 11.5 g/l sodium citrate dehydrate and 2% glucose, 0.04 g/l Adenine, 0.08 g/l Uridine). For persistent and non-persistent clinical isolates, cells were grown overnight in YPD and 10^5^ cells were spread onto 15 ml YPD plates. To facilitate comparisons between casitone and YPD disk diffusion assays, a subset of control strains with different FoG levels were included in both types of assays (SC5314, GC75, P87). The fraction of growth and radius of inhibition levels, referred to as FoG and RAD throughout the manuscript, represent parameters measured at 20% drug inhibition (FoG_20_ and RAD_20_, respectively). For disk diffusion assays performed with auxotrophic strains, supplementary amino acids were added to the media (0.04 g/l Histidine and 0.04 g/l Arginine). For disk assays with drug adjuvants, the following concentrations of drugs were used: 0.5 μg/ml geldanamycin, 0.5 μg/ml radicicol, 0.5 μg/ml FK506, 0.4 μg/ml cyclosporine A, 20 μg/ml fluoxetine, 5 ng/ml aureobasidin A, 0.5 ng/ml rapamycin, 10 μg/ml fluphenazine, 50 μg/ml doxycycline, 12.5 ng/ml staurosporine and 0.25 μg/ml Tunicamycin. A single 25 μg FLC disk (6 mm, Oxoid, UK or Liofilchem, Italy) was placed in the center of each plate, plates were then incubated at 30°C for 48 h, and each plate was photographed individually. Analysis of the disk diffusion assay was done using the *diskImageR* pipeline^65^ and the R script is available at https://github.com/acgerstein/diskImageR/blob/master/inst/walkthrough.R. Several controls indicated that drug in the disk is stable: incubation of the disk on the plate for 24 h prior to plating cells did not change FoG levels. In addition, incubation of the disk on the plate for 24 h prior to plating cells did not affect FoG levels, neither did addition of a fresh disk to the plate after 24 h.

### ScanLag assay

The ScanLag assay was adapted from^74^ with minor modifications. Strains were streaked from glycerol stocks onto YPD agar and incubated for 24 h at 30°C. Colonies were suspended in 1 ml PBS, diluted to 10^4^ cell/ml and ∼ 500 were spread onto casitone plates with or without 10 μg/ml FLC (Sigma-Aldrich, St. Louis, MO). Plates were placed on the scanners at 30°C and scanned every 30 min for 96 h. Image analysis was done in MATLAB using the “ScanLag” script^74^ that was adapted for yeast cells by changing the identification of the size of the colony to a minimum of 20 pixels.

### Viability assays

Replica plating was performed from disk diffusion plates that were incubated at 30°C for 48 h. Master plates were inverted and pressed firmly on a sterile cotton velveteen stamp and then transferred to new casitone plates containing no drug. Replica plates were incubated at 30°C for 48 h, and then each plate was photographed individually.

### Enzyme-linked immunosorbent assay (ELISA)

The ELISA protocol was modified from^131^. Briefly, *C. albicans* cells were grown for 6 h at 30°C with shaking (220 RPM) in YPD with or without 10 µg/ml FLC (Sigma-Aldrich, St. Louis, MO) and harvested at exponential phase (OD_600_ 0.6–0.8). Cells were centrifuged for 10 min (3000 rpm, 4°C), and washed once with ice-cold PBS. Pellets were resuspended in 200 μl lysis buffer (50 mM Hepes pH 7.5, 150 mM NaCl, 5 mM EDTA, 1% Triton X100, protease inhibitor cocktail (Roche Diagnostics)) together with acid-washed glass beads. Cells were then mechanically disrupted by vortexing for 30 min at 4°C. Cell lysates were diluted 1:10 in PBS and subjected to BCA analysis (Pierce Biotechnology, Rockford, IL) to determine protein concentrations. Cell lysates at 10 µg/ml concentration were incubated in 96-well ELISA plates for 18 h at 4°C. Wells were washed with PBST (PBS +0.05% Tween-20), blocked with 1% skim milk in PBST for 2 h at 37°C and washed. Rabbit polyclonal anti Hsp90 antibody (Dundee Cell Products, Scotland) was diluted 1:1000 in blocking buffer applied overnight at 4°C. Horseradish peroxidase-coupled donkey anti-rabbit IgG (1:1000 dilution, Jackson ImmunoResearch Laboratories, West Grove, PA) was incubated for 1 h at 37°C followed by washing. Detection was done with TMB (3,3’,5,5’-tetramethybenzidine). The reaction was terminated with 0.16 M sulfuric acid and absorbance was measured at 450 nm in an ELISA plate reader.

### FLC-Cy5 uptake measured by flow cytometry

*C. albicans* cells were grown overnight in YPD media at 30°C Cultures where diluted 1:100 in 3 ml casitone medium and incubated for 3 h at 30°. Drugs were added to a final concentration of 10 μg/ml for FLC, and 1 μg/ml for FLC-Cy5^81^. Cells were harvested every 30 min and diluted 1:4 in 50% TE (50 mM Tris pH 8:50 mM EDTA). Data was collected from 25,000-35,000 cells per time point using 561 nM EX and 661/20 nM EM in a MACSQuant flow cytometer and gated to by SSC<10^3^ A.U and FSC>10^4^ A.U in (SSC vs FSC) to eliminate small debris particles. Analysis was done using FlowJo 8.7 software.

### Efflux of Rhodamine 6G (R6G) as measurement of drug efflux capacity

The assay was adapted from^132^ with minor modifications. In brief, strains were streaked from glycerol stock onto YPD plates and incubated for 24 h at 30°C. Colonies were resuspended in 3 ml casitone medium and grown overnight in 30°C. Cultures where diluted 1:100 in 5 ml casitone medium and incubated for 3 h at 30°. Cells were centrifuged, washed in 5 ml PBS (pH 7), and resuspended in 2 ml PBS with or without 10 μg/ml FLC. Cells suspensions were incubated for 1 h at 30°C and R6G (Sigma-Aldrich, St. Louis, MO USA) was added at 10 μg/ml to allow R6G accumulation (1 h). Next, cells were washed twice with PBS at 4°C, and resuspended in a final volume of 300 μl PBS. 50 μl of individual suspensions were diluted in 150 μl PBS and aliquoted into a 96-well microtiter plate. Baseline emission of fluorescence (excitation 344 nm, emission 555 nm) was recorded for 5 min in a Tecan plate reader at 30°C (Infinite F200 PRO, Tecan, Switzerland), in relative fluorescence units (RFU), and 1% D-glucose was next added to each strain to initiate R6G efflux. Negative controls contained no glucose and data points were recorded for 90 min in triplicates at 1 min intervals.

### *G. mellonella* virulence assays

*G. mellonella* larvae were obtained from Vanderhorst Wholesale (Saint Marys, OH) and the infection protocol was adapted from Li *et al*^133^. Larvae were kept at 15**°**C and used within 1 week of delivery. Larvae were randomly selected for each experiment in groups of 12 and those showing signs of melanization were excluded. *C. albicans* inoculums were prepared from cultures grown overnight at 30**°**C in YPD and washed twice with sterile PBS. Cells were counted using a hemocytometer and adjusted to 3 × 10^7^ cells/ml in sterile PBS. Each larva was initially injected with 3 × 10^5^ cells via the last left pro-leg using a 10 μl glass syringe and a 26S gauge needle (Hamilton, 80300). A second injection with either FLC (1 mg/kg), FLC + fluphenazine (FNZ, 10 mg/kg), or sterile PBS was done via the last right pro-leg 1.5-2 h post infection. The inoculum size was confirmed by plating of fungal cells on YPD. Infected larvae and PBS injected controls were maintained at 37**°**C for 14 days and checked daily. Larvae were recorded as dead if no movement was observed upon contact. Virulence assays were performed with strain SC5314, with SC5314-derived isolates passaged for 12 days in either YPD alone (2 ‘low’ FoG isolates) or YPD plus 1 μg/ml FLC (4 ‘high’ FoG isolates), and with isolates NP03 and S12-01 from a clinical isolate set (see Supplementary Table 1). Experiments were performed in duplicate (n = 24 larvae) and statistical differences between larval groups were tested using the Mantel-Cox test. Negative controls included PBS, FLC alone or FLC + FNZ for groups of 24 larvae. No significant killing of larvae was observed in either of these conditions.

### Growth assays

To determine growth parameters (doubling time and lag phase duration), *C. albicans* cells were seeded at a concentration of 2 × 10^5^ cells/ml into 96-well plates containing YPD media. Plates were incubated for 48 h at 30°C with continuous shaking in a Tecan plate reader (BioTek) and the optical density reading (OD_600_) was recorded every 15 min. Measurements of doubling time and lag phase duration were determined using BioTek Gen5 software and from 3 biological replicates, each performed in duplicate.

### Microcolony assays

Microcolony assays were adapted from^72^ with minor modifications. Colonies were resuspended in 3 ml casitone medium and grown overnight in 30°C. Cultures were diluted 1:30 in casitone medium and incubated at 30°C until cultures reached logarithmic phase (∼3-4 h) and diluted to 10^4^ cells/ml in casitone medium. For microscopy, glass-bottomed 24-well plates (De-GROOT, Israel) were coated with 1 ml of a 200 μg/ml Concanvalin A solution (Type VI, Sigma-Aldrich, St. Louis, MO USA) for 3-4 h. Wells were washed twice with 1 ml ddH20 and 10^4^ cells/ml in a 2 ml volume were added to wells with or without 10 μg/ml FLC. Plates were sealed with an optically clear film (Axygen, Corning, Israel) and spun at 1200 rpm for 1 min. Images where captured in 100 fields per well for 48 h in 1 h intervals using a Nikon Ti Eclipse microscope equipped with a full-stage environmental chamber (set to 30°C) with a Nikon Plan Apo 10x (0.45 numerical aperture) objective using Nikon Elements AR software. The focusing routine utilized manual assignment for each well based on a single field^134^. Custom software for conducting microcolony experiments, including computing and tracking microcolony areas over time was adapted from^72^. Specific growth rates were calculated for 1500-5000 colonies per strain^72^ by regressing the natural log of microcolony area over time. Growth rates were calculated separately for two time intervals (0-5 h and 10-15 h) and required to have an R^2^>0.9. Replicate wells (3 per condition) were consistent and growth rate distributions were pooled.

### Statistical analyses

All experiments represent the average of two or more biological replicates, with two technical replicates of each. Error bars represent the standard deviation. Statistical analyses were performed using two-tailed Student’s t tests, one way ANOVAs and Tukey’s multiple comparison tests using Microsoft Excel 2016 (Microsoft) and Prism 6 (GraphPad); R^2^ tests were used for linear regressions. Significance was assigned for *P* values smaller than 0.05, asterisks denote *P* values as follows: ***, *P* < 0.001; **, *P* < 0.01; *, *P* < 0.05.

## Data availability

The authors declare that all data supporting the findings of the study are available in this article and its Supplementary Information files, or from the corresponding author upon request.

## Acknowledgements

We thank members of the Berman and Bennett labs for stimulating discussions throughout the work. We are grateful to G. Palmer, M. Lohse, O. Homann, A. Johnson, R. Ben-Ami, T. White, D. Soll, D. McCallum, F. Odds, D. Sanglard, L. Cowen, P.T. Magee, B. Cormack, P. Magwene, M. Kupiec, W. Fonzi, S. Noble, B.M. Vincent, S. Lindquist, and A. Colombo for providing strains and P. Lipke, M. Fridman, R. BenAmi, N. Dror, A. Selmecki, A.C. Gerstein and A. Forche for helpful comments on the manuscript. This work was funded by European Research Council Advanced Award 340087 (RAPLODAPT) to JB, by Fundação de Amparo a Pesquisa do Estado de São Paulo (FAPESP, Grant 2017/02203-7), Brazil to ALC, by National Institutes of Health grant AI081704 to RJB and by a Brazil Initiative Collaborative Grant from the Watson Institute to RJB and IVE.

## Author contributions

IVE, RJB and JB designed the study; AR, IVE and AD collected the data; AR and IVE analyzed the data, MB produced and analyzed data in Fig. 3a,d, SZ performed the experiment in Supplementary Fig. 7g, ESS constructed mutant strains, NZ implemented and helped analyze data in Fig. 2b, ALC provided clinical isolates and clinical input to the paper; AR, IVE, RJB and JB wrote the manuscript.

